# Evolution of phenotypic variance in response to a novel hot environment

**DOI:** 10.1101/2021.01.19.427270

**Authors:** Wei-Yun Lai, Christian Schlötterer

## Abstract

Shifts in trait means are widely considered as evidence for adaptive responses, but the impact on phenotypic variance remains largely unexplored. Classic quantitative genetics provides a theoretical framework to predict how selection on phenotypic mean affects the variance. In addition to this indirect effect, it is also possible that the variance of the trait is the direct target of selection, but experimentally characterized cases are rare. Here, we studied gene expression variance of *Drosophila simulans* males before and after 100 generations of adaptation to a novel hot laboratory environment. In each of the two independently evolved populations, the variance of 125 and 97 genes was significantly reduced. We propose that the drastic loss in environmental complexity from nature to the lab may have triggered selection for reduced variance. Our observation that selection could drive changes in the variance of gene expression could have important implications for studies of adaptation processes in natural and experimental populations.

## Background

Most studies of adaptation rely on shifts in trait mean as signal of selective response (Jakšić et al., 2020; Lemos, Bettencourt, Meiklejohn, & Hartl, 2005; Mallard, Nolte, Tobler, Kapun, & Schlötterer, 2018; Nuzhdin, Wayne, Harmon, & McIntyre, 2004; Oleksiak, Churchill, & Crawford, 2002; Whitehead & Crawford, 2006). The variance of the trait in a population, which is the prerequisite for an adaptive response (Bull, 1987; Falconer & Mackay, 1963), has received considerably less attention. As a result, our understanding of the evolution of phenotypic variance is still rather limited. Probably, most progress has been made in quantitative genetics, describing the evolution of phenotypic variance in response to a sudden shift in trait optimum (Bulmer, 1972; Chevalet, 1994; Kimura & Crow, 1964; Turelli, 1984). For large populations and traits controlled by many unlinked loci with equal effect, changes in trait optimum are not expected to affect the phenotypic variance (Hayward & Sella, 2019; Lande, 1976). In contrast, a much more complex picture is expected when the effect sizes are not equal, the population size is finite, or the traits have a simpler genetic basis (Barton & Keightley, 2002; Barton & Turelli, 1987; Franssen, Kofler, & Schlötterer, 2017; Jain & Stephan, 2015; Keightley & Hill, 1989).

In addition to these indirect effects on phenotypic variance, it is also possible that it is not the trait mean, but rather the variance of the trait that is the target of selection. For instance, stabilizing selection may reduce the variance of a trait (Schmalhausen, Isadore, & Dobzhansy, 1951). Canalization, one potential consequence of stabilizing selection (Le Rouzic, Álvarez-Castro, & Hansen, 2013), describes the phenomenon that genetic and environmental perturbations can be buffered and henceforth reduce the phenotypic variance. A classic textbook example for a canalization factor is the heat shock protein Hsp70. Mutations of this chaperone gene result in increased phenotypic variance due to the unmasking of genetic variation (Rutherford & Lindquist, 1998). Because canalization differs between populations, it has been proposed that it may also evolve (Flatt, 2005; Rice, 1998).

The selective forces operating after a shift in habitat can be characterized by the evolution of gene expression mean and variance using an experimental evolution setup. When stabilizing selection becomes stronger, even with no directional selection being involved, we expect a pronounced reduction in variance. When directional selection comes into place, for genes/traits with simple genetic architectures, we expect to observe associated changes of both mean and variance. In contrast, only mean evolution is expected when directional selection is operated on polygenic traits/gene expression. We studied gene expression variance of *Drosophila simulans* males before and after 100 generations of adaptation to a novel hot laboratory environment. We found no detectable difference in variance evolution between the genes with significant changes in mean expression and those without, and thus discussed the polygenic basis of the expression evolution for most genes in a separate manuscript (Lai, Nolte, Jaksic, & Schlötterer, 2021). In this study, we focused on a small subset of genes, which changed significantly their phenotypic variance during 100 generations of adaptation. Genes which evolved a reduced expression variance were strongly enriched in the gut and food processing. We discuss how a potential increase in stabilizing selection pressure could reduce gene expression variance after a drastic change in environmental complexity from nature to the lab. Our results suggest that the shift in environment can drive the evolution of phenotypic variance, which requires further attention in the interpretation of phenotypic evolution.

## Materials and methods

### Experimental evolution

The setup of populations and evolution experiment have been described by (Barghi et al., 2019). Briefly, ten outbred populations seeded from 202 isofemale lines were exposed to a laboratory experiment on standard *Drosophila* medium (300g Agar + 990g sugar beet syrup + 1000g Malt syrup + 2310g corn flour + 390g soy flour +900g yeast in 37.5L water) at 28/18 °C with 12hr light/12hr dark photoperiod for more than 100 generations. Each population consisted of 1000 to 1250 adults at each generation.

### Common garden experiment

The collection of samples from the evolution experiment for RNA-Seq was preceded by two generations of common garden (CGE) to control for transgenerational and environmental effects. Most phenotypic differences observed in a common garden experiment should reflect genetic differences. The common garden experiment was performed at generation 103 of the evolution in the hot environment and this CGE has been described in (Barghi et al., 2019; Hsu et al., 2019, 2020; Jakšić et al., 2020). In brief, an ancestral population was reconstituted by pooling five mated females from 184 founder isofemale lines. Each isofemale line was maintained at small population size (typically less than 50 individuals) at 18 °C for ∼50 generations before the reconstitution on standard laboratory food. Potential adaptation to the lab environment with the residual heterogeneity or de novo mutation in each line could be possible. Nevertheless, as discussed (Barghi et al., 2019), given the small effective population size during the maintenance of each line, most variants/mutations are effectively neutral and thus adaptation is unexpected. This has also been experimentally tested by contrasting allele frequencies in populations which were reconstituted shortly after the establishment of the lines and after 50 generations, but no significant allele frequency differences were found. This supports the idea that no consistent adaptation pattern can be recognized in isofemale lines maintained in the laboratory (Nouhaud, Tobler, Nolte, & Schlötterer, 2016). Furthermore, it is important to keep in mind that selection on standing genetic variation could only occur until an isofemale line is inbred. The low Ne may, however, allow the accumulation of deleterious mutations within the isofemale lines. Nevertheless, since most mutations occurring in the laboratory are mostly recessive (unlike mutations spreading in natural populations) (Charlesworth & Charlesworth, 2010), we expect very little impact as the two generations of common garden resulted in heterozygous flies, where recessive deleterious mutations are masked. The reconstituted ancestral populations and two independently evolved populations at generation 103 were reared for two generations with egg-density control (400 eggs/bottle) at the same temperature regime as in the evolution experiment. Freshly eclosed flies were transferred onto new food for mating. Sexes were separated under CO_2_ anesthesia at day 3 after eclosure, left to recover from CO_2_ for two days, and at the age of five days, whole-body mated flies of each sex were snap-frozen at 2pm in liquid nitrogen and stored at -80°C until RNA extraction. In this study, more than 30 individual male flies from two reconstituted ancestral populations (no. 27 and no. 28) and two randomly selected evolved populations (no. 4 and no. 9) were subjected to RNA-Seq. While it would have been desirable to sequence all 10 available populations, due to budget constraints, we had to limit our analysis to two populations, since this already involved the generation and sequencing of more than 120 independent libraries.

### RNA extraction and library preparation

Whole bodies of individual male flies were removed from the -80°C freezer and immediately homogenized in Qiazol lysis reagent (Qiagen, Hilden, Germany). The homogenate was treated with DNase I followed by addition of chloroform, centrifugation and mixture of the upper phase with 70% ethanol as described for the Qiagen RNeasy Universal Plus Mini Kit. The mixture was subsequently loaded onto a RNeasy MinElute Spin column as provided by the RNeasy Plus Micro Kit (Qiagen, Hilden, Germany), and all washing steps were performed according to the instructions for that kit. All resulting total RNA was used to prepare stranded mRNA libraries on the Neoprep Library Prep System (Illumina, San Diego, USA) following the manufacturer’s protocol: Neoprep runs were performed using software version 1.1.0.8 and protocol version 1.1.7.6 with default settings for 15 PCR cycles and an insert size of 200bp. All libraries for individuals of ancestral population no. 27 and evolved population no. 4 were prepared with library cards of lot no. 20180170; all libraries for individuals of ancestral population no. 28 and evolved population no. 9 were prepared with library cards of lot no. 20178099. 50bp single-end reads were sequenced on an Illumina HiSeq 2500.

### RNA-Seq data processing and quality check

All RNA-Seq reads were trimmed using ReadTools (Gómez-Sánchez and Schlötterer, 2018) with quality score of 20 and aligned to *Drosophila simulans* reference genome (Palmieri, Nolte, Chen, & Schlötterer, 2015) using GSNAP (Wu, Reeder, Lawrence, Becker, & Brauer, 2016) with parameter setting -k 15 -N 1 -m 0.08. The reference genome has been annotated using the flybase gene IDs of *D. melanogaster* for all orthologous genes (Palmieri et al., 2015). Exon-aligned reads were piped into Rsubread (Liao, Smyth, & Shi, 2019) to calculate read counts of each gene, and raw read counts of each gene were normalized for library size and subjected to the TMM normalization in edgeR (Robinson, McCarthy, & Smyth, 2010). Samples with severe 3’-bias were removed based on the gene-body coverage plot (Jakšić & Schlötterer, 2016; Wang, Wang, & Li, 2012). Additionally, we removed five individuals where the freezing process may have led to detachment of body parts, such as eyes or heads. Specifically, we compared gene expression between each sample and all other samples and identified the samples exhibiting outlying expression pattern. Tissue enrichment analysis was performed for genes with at least 2-fold lower expression in the outlying samples to provide significant evidence of tissue detachment. After quality control, each population remained approximately 20 individuals (Supplementary file 1). Only genes with at least 1 count per million (CPM) were included in the analyses to avoid extremely lowly expressed genes.

### RNA-Seq data analysis

For all RNA-Seq data, we only compared samples which were prepared with library cards from the same lot number to avoid batch effects (Comparison 1: evolved population no. 4 vs. reconstituted ancestral population no. 27; Comparison 2: evolved population no. 9 vs. reconstituted ancestral population no. 28).

For the analysis of the evolution of mean/variance in gene expression, we applied natural log transformation (Heath, Bulfield, Thompson, & Keightley, 1995) to eliminate the strong positive mean-variance correlation in RNA-Seq data due to the nature of the negative binomial distribution (Supplementary figure 1) (Lai et al., 2021). The mean and variance of the expression of each gene (lnCPM) were estimated in each population. The change of gene expression mean was determined with the linear modeling framework implemented in limma (Ritchie et al., 2015). The change of gene expression variance was determined with the F test between the variance within the ancestral population and the variance within the evolved population for each gene. P-value adjustment was performed using the Benjamini-Hochberg false discovery rate (FDR) correction. To demonstrate concordant evolution in both replicates, we tested whether more genes with significant variance/mean evolution in the same direction were detected than expected by chance with Fisher’s exact tests. In addition, we investigated to which extent the heterogeneous response can be explained by stochastic variation due to the experimental noise or sampling processes. Within the first comparison, we resampled twice 15 samples from both the evolved and reconstituted ancestral populations, performed two tests for variance (contrasting the evolved and ancestral samples) and estimated the consistency between the tests using the Jaccard Index as 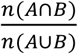, where A and B are the candidate gene sets of the two tests. This process is repeated 100 times, generating the null distribution of Jaccard Indices considering sole stochastic effects within an evolutionary replicate. We applied a similar sampling scheme of 15 individuals to evaluate the consistency in evolution between populations. We contrasted the evolved and ancestral samples for the two populations (rather than for two samples of the same population, which was done to obtain the null distribution). The observed Jaccard indices from 100 times of resampling were compared to the null distribution.

We compared the proportion of genes with significant differences in mean in genes with and without significant reduction in variance to evaluate whether the reduced variance is associated with the selection on mean. Furthermore, we correlated the changes in gene expression variance (ln(F)) with the magnitude of mean evolution (absolute value of mean difference) for the genes with significant changes in either expression mean or variance.

### Gene ontology and tissue enrichment analysis

We used ClueGO (Bindea et al., 2009) to perform gene ontology (GO) enrichment analyses of the candidate genes have significant change on variance. To understand in which tissues the genes of interest are expressed, we made use of tissue-specific expression profiles of adult males of *Drosophila melanogaster* on flyatlas2 (Leader, Krause, Pandit, Davies, & Dow, 2018). This data set includes 13 tissues in male flies. Genes that are expressed 4-fold higher in a given tissue than in the whole body are identified. Fisher’s exact test was performed to test if the genes of interest are enriched for genes highly expressed in one tissue. P-value adjustment was performed using the Benjamini-Hochberg false discovery rate (FDR) correction.

### Microbiome diversity in ancestral and evolved populations

To explore the heterogeneity in gut microbiome, we performed 16S-rRNA amplicon sequencing on three remaining individual males of the ancestral and evolved populations from the same common garden experiment (Supplementary file 1).

We used primers designed to amplify the V3-V4 hypervariable regions of the 16S rRNA gene. The primers had an overhang to match Nextera Index primers (Forward primer: 5’- TCGTCGGCAGCGTCAGATGTGTATAAGAGACAG-CCTACGGGNGGCWGCAG-3’, Reverse primer: 5’-GTCTCGTGGGCTCGGAGATGTGTATAAGAGACAG-GACTACHVGGGTATCTAATCC-3’). PCR products were amplified with 30 cycles at an annealing temperature of 65°C, purified using AMPure XP beads (Beckman Coulter, Carlsbad, CA) and subjected to a second PCR to introduce dual index sequences using Nextera XT Index Kits (Illumina, San Diego, CA). In the second PCR, we used 6 cycles and an annealing temperature of 55°C, and both PCRs were carried out in 5μl total volume using the NEBNext Ultra II Q5 Mastermix (New England Biolabs, Ipswich, MA). The final amplicons were again purified, quantified using the Qubit HS assay kit (Invitrogen, Carlsbad, CA), and 125 bp paired-end reads were sequenced on an Illumina HiSeq 2500.

The 16S-rRNA sequence data were trimmed using ReadTools (Gómez-Sánchez & Schlötterer, 2018) with quality score of 20. Unpaired reads were removed. Owing to the variation in sequencing depths between samples, all samples were down-sampled to the lowest depth. Each bam file was converted into a fastq.gz file and analyzed with Kraken2 (Wood, Lu, & Langmead, 2019) following the recommended parameters and the estimation of genus abundance was corrected by Bracken (Lu, Breitwieser, Thielen, & Salzberg, 2017).

Genus abundance of the microbiome community in each sample was obtained. With the filtration (read number < 5), extremely lowly abundant genera were excluded. β-diversity (Tuomisto, 2010) was then calculated to evaluate the heterogeneity of the microbiome complexity among the three individuals from the same population.

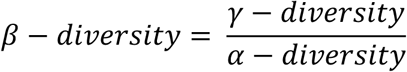

where γ-diversity is the genera species richness in a population and α-diversity is the mean richness within an individual.

### Simulation study

We performed forward simulations with MimicrEE2 (Vlachos & Kofler, 2018) using the *qff* mode to illustrate the influence of the genetic architecture on the evolution of phenotypic variance during the adaptation to a new trait optimum. With 189 founder haplotypes (Barghi et al., 2019), we simulated quantitative traits under the control of 20 numbers of loci with an effective population size of 300. For each trait, we assume an additive model and the negative correlation (r=-0.7) between the effect size (*α*∼Γ(100,15)) and starting frequency (Barghi et al., 2019). We used *correlate()* function implemented in “fabricatr” R package (Blair et al., 2019) to generate the effect sizes with negative correlation (r = -0.7) with starting frequency. The sum of effect sizes of each trait was normalized to 1. We assumed heritability h^2^ = 0.6. To simulate strong stabilizing selection without trait optimum shift, we provided the fitness function: 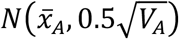, where 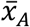 is the ancestral phenotypic mean and *V*_*A*_ is the ancestral genetic variance. For the neutrality case, we assumed the same fitness for each individual. For each trait under each scenario, the phenotypic variance was calculated at different generations and normalized to the ancestral phenotypic variance at generation 1 to investigate the dynamic of phenotypic variance during the evolution.

### Transcription factor enrichment analysis

Transcription factor enrichment analysis among the genes with significant decreased variance in the midgut was done with Rcistarget (version 1.0.2) (Aibar et al., 2017). First, enrichment of cis-regulatory elements (CREs), 5kb upstream and intronic sequences, of the genes of interest was identified. The motif-search database used here was based on the latest motif ranking files of *Drosophila* species (“dm6-5kb-upstream-full-tx-11species.mc8nr.feather”). Parameter setting used in this analysis is as following: nesThreshold = 5 and aucMaxRank = 0.05. The predicted transcription factors (TFs) were considered as candidate TFs regulating the genes of interest.

## Results

### Rapid changes in gene expression variance during adaptation

We measured the gene expression of 19-22 whole body male individuals from two populations independently evolved in a novel hot environment and reconstituted ancestral populations (81 samples in total). After adapting for 100 generations to the high temperature regime, the transcriptomic response of hot-evolved populations was significantly diverged from their ancestors. Principal Component Analysis indicated that PC1 explained 11.9% of the total variation and separated the hot-evolved flies from their ancestor which reflects the clear adaptive signatures to the novel, hot temperature regime (Lai et al., 2021). The variances of the expression of each of the 10,583 genes were estimated and compared between the reconstituted ancestral populations and the two evolved populations. Only samples from the same lot number of library preparation (Supplementary file 1) were compared (See Materials and Methods).

In both hot evolved populations, a small number of genes (166 and 148) significantly changed in expression variance after 100 generations of adaptation (F-test, FDR < 0.05; Figure 1; Supplementary file 2). Among the 166 genes with a significant change in variance in population 1, the variance of 125 genes decreased while only 41 genes showed a variance increase. This is a significant difference in the directionality of phenotypic variance evolution (χ^2^=42.51, p-value < 7.0×10^−11^). A similar difference was seen in population 2 (97 decreased and 51 increased; χ^2^=14.30, p-value < 1.6×10^−4^). 14 genes with a significant variance change in the same direction (11 decreased and three increased) were shared between the two independently evolved populations, which is significantly more than expected by chance (Fisher’s Exact Test, odds ratio = 7.07, p-value < 7.52 × 10^−8^). Hence, the same selection pressure was likely to drive the loss of expression variance in the two populations evolving independently in the same novel hot environment. This parallel variance change persisted when we lowered the significance threshold of the F-test to FDR < 0.1 that yield 274 and 229 genes with significant changes in variance in both populations (Fisher’s Exact Test, odds ratio = 4.49, p-value < 2.49 × 10^−8^).

**Figure 1.**
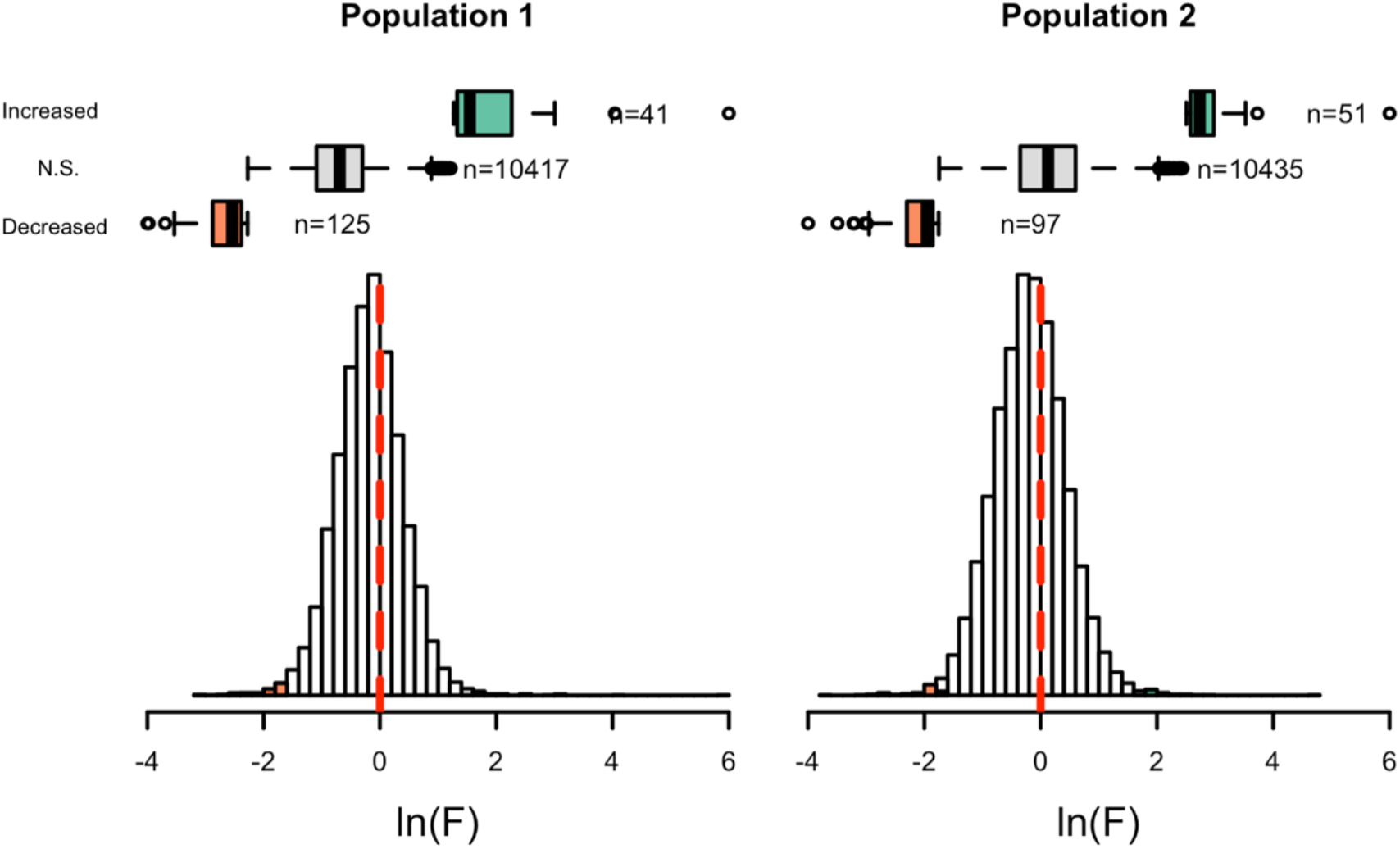
Evolution of gene expression variance. The distribution of the change in gene expression variances (ln(F)) during the evolution experiment in the 1^st^ (left panel) and 2^nd^ (right panel) population. Boxes in salmon indicate the genes with decreased variance in both populations (n=125 and 97) and boxes in green represent genes with increased variance (n=41 and 51). Boxes in grey include the other genes without significant change in variance (n=10417 and 10435).

### Reduction in expression variance is not associated with changes in mean expression levels

Previous analyses of the same evolved populations identified many genes with a significant shift in mean expression and associated biological processes in response to adaptation to the new temperature regime in the laboratory (Barghi et al., 2019; Hsu et al., 2020; Jakšić et al., 2020). Consistent with these results, the comparison of ancestral and evolved populations in this study identified 2,775 genes in the first replicate and 2,677 genes in the second replicate which significantly changed mean expression (Supplementary file 3). Most of them were shared with those detected in previous studies (Jakšić et al., 2020) (Fisher’s exact test, odds ratio = 4.1 and 3.0, FDR < 2.2 × 10^−16^). 94.1% of the genes with a significant mean expression change in both evolved populations changed in the same direction, significantly more than expected by chance (χ^2^ = 898.03, p-value < 2.2 × 10^−16^). This concordance suggests that most of the altered expression means are mainly driven by selection, rather than by drift. If these changes in mean expression are driven by selection either on *cis*-regulatory variation or few *trans*-acting factors, it is expected that changes in mean are also associated with reduced expression variance. For genes with a significant decrease in variance, the proportion of genes with significant change in mean was slightly higher compared to genes without variance change, but the difference is not significant (Figure 2, χ^2^ = 3.96 *and* 1.96, FDR > 0.05). Further support for the independence of variance and mean changes in expression comes from the observation that, out of 11 genes with reduced variance in both populations, only one gene showed a significant change in mean in both populations. Furthermore, the magnitude of mean changes was basically not correlated with the changes in variance among genes with significantly altered means or variances (Supplementary figure 2, Spearman’s rho = 0.047 and -0.001). These results suggest that mean and variance are evolving independently, probably because of different selection pressures.

**Figure 2.**
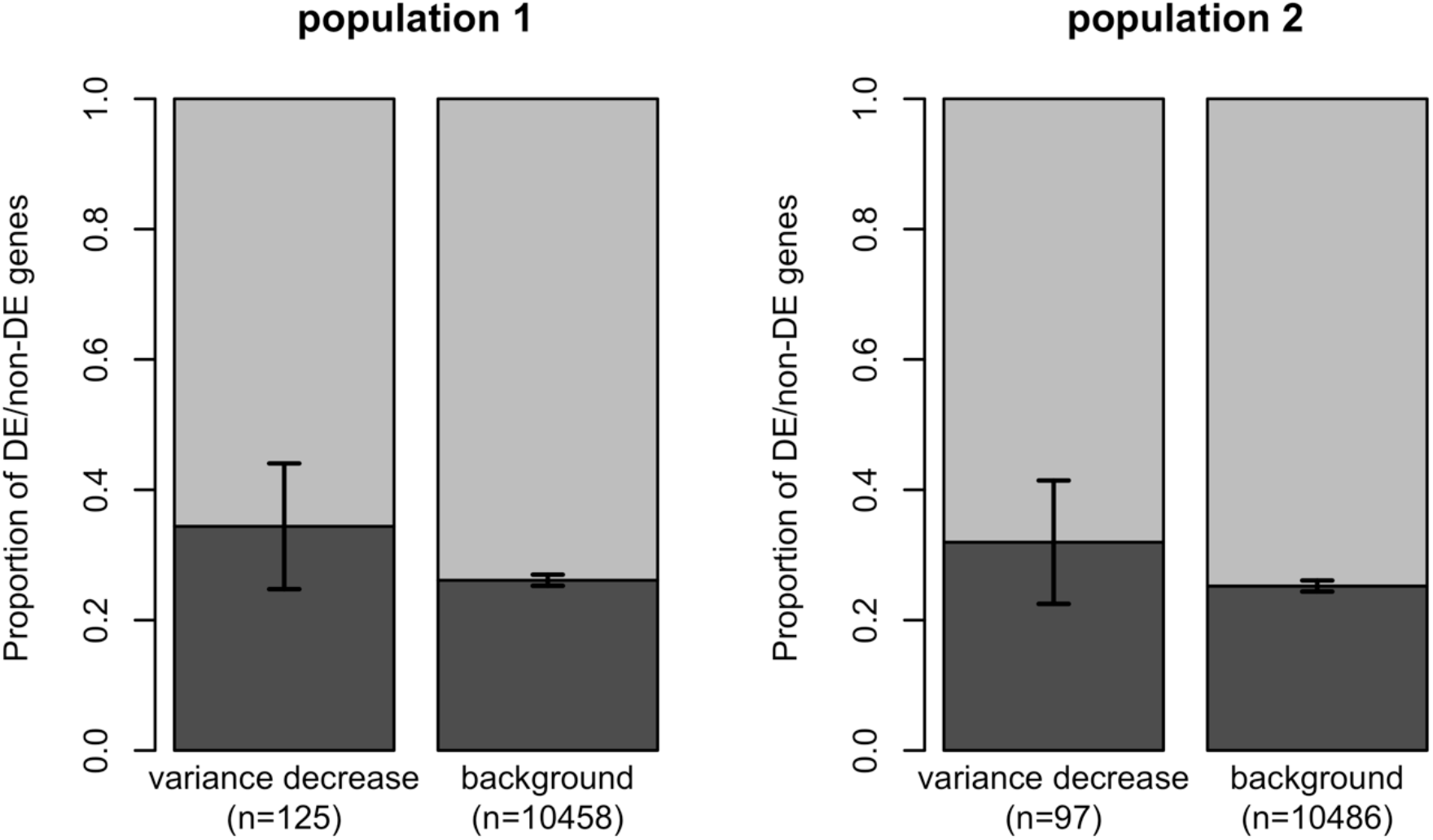
No significant enrichment of DE genes for the genes with significant variance change. Despite a trend for a slightly higher fraction of genes with a change in expression mean among genes with a significant change in expression variance, considering both populations jointly, no significant difference was noticed compared to genes without significant variance decrease (χ^2^ = 3.96 *and* 1.96, FDR > 0.05). The y-axis denotes the proportion of DE (dark grey)/non-DE (light grey) genes in the genes with significant variance decrease and without (background) for two populations. The error bar indicates the 95% confidence interval.

### Digestive genes in midgut rapidly decreased their transcriptional variance

In order to characterize processes that could explain the significant changes in gene expression variance, we searched for gene ontology (GO) or tissue-specific expression enrichment. In both populations, genes with increased variance had no consistent enrichment in any biological processes or tissue-specific expression (Supplementary file 4 and 5). In contrast, for the genes with decreased variance, the enrichment for expression in the midgut was significant for both populations (Fisher’s exact test, FDR < 0.05, Figure 3, Supplementary file 4). We note that allometric changes during adaptation cannot explain this pattern since all genes expressed in the affected tissues would be modified. We found, however, that different sets of midgut-expressed genes changed their expression variance in the two populations (Supplementary file 6). GO enrichment analysis identified also catabolism-related processes (e.g.: “organic substance catabolic process”, “carbohydrate metabolic process” and “organonitrogen compound catabolic process”) in both populations (Supplementary file 5). In addition to the consistent enrichment in the midgut and catabolic processes, we also observed an enrichment for digestive enzymes (Lemaitre & Miguel-Aliaga, 2013) (Fisher’s exact test, odds ratio = 4.21 and 3.53, FDR < 0.01, Supplementary file 6), indicating that different sets of digestive genes in midgut rapidly decreased their transcriptional variance in the two populations during 100 generations of adaptation. The enrichment in midgut and digestive genes persisted when we lowered the significance threshold of the F-test (FDR < 0.1, supplementary file 4), indicating that our result does not depend on a specific cutoff to define the genes with reduced gene expression variance.

**Figure 3.**
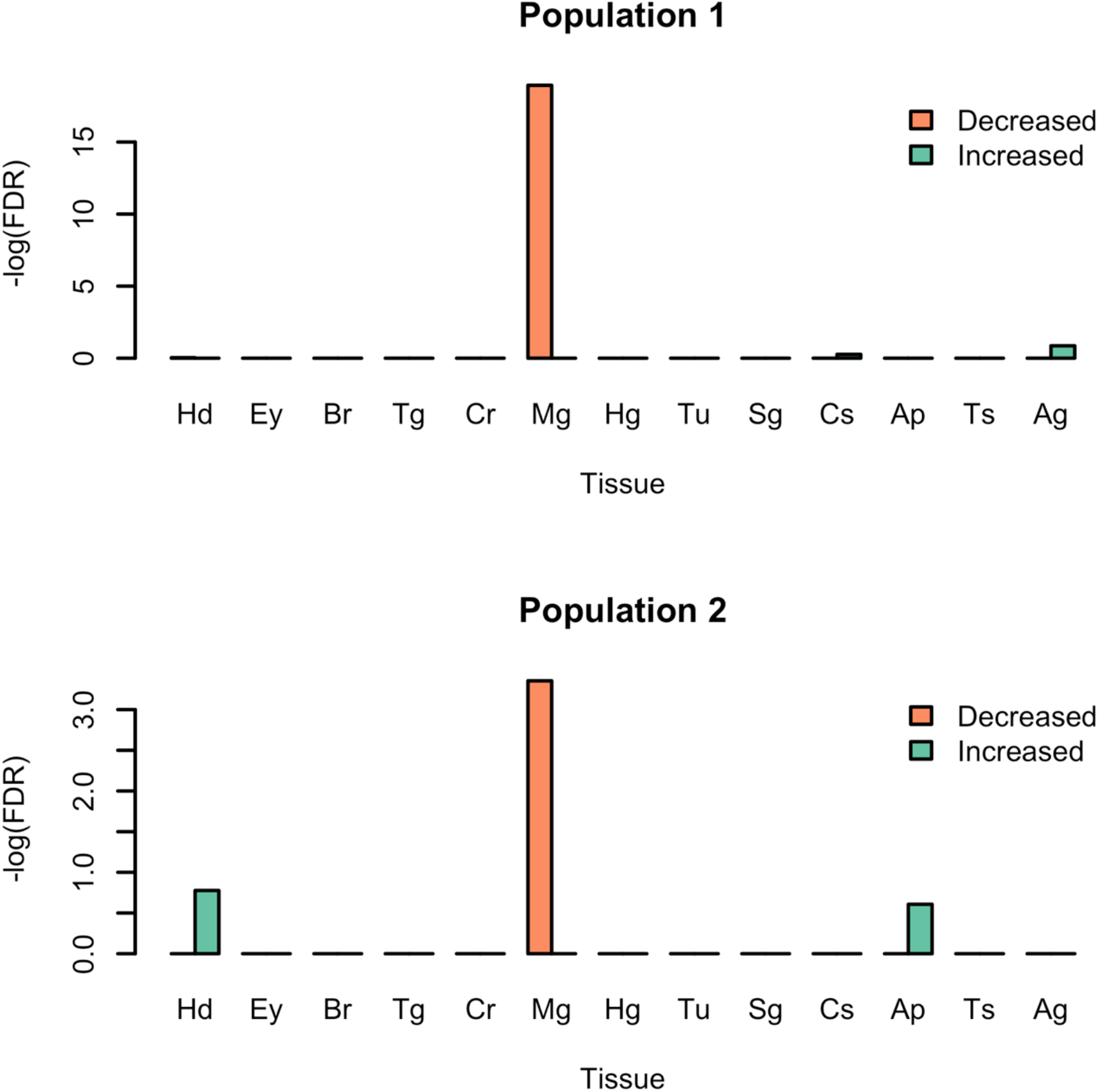
Tissue enrichment of genes with significant changes in expression variance. The bars indicates the significance (-ln(FDR)) of enrichment for genes with significant variance changes (orange: genes with decreased variance; green: genes with increased variance) among genes with tissue-specific gene expression pattern (Hd-head, Ey-eye, Br-brain, Tg-thoracoabdominal ganglion, Cr-crop, Mg-midgut, Hg-hindgut, Tu-malpighian tubule, Sg-salivary gland, Cs-carcass, Ap-rectal pad, Ts-testis and Ag-accessory glands). In both populations, a significant enrichment (Fisher’s exact test, FDR < 0.001) can be found in the midgut for genes with reduced expression variance.

Next, we focused on the 11 genes that showed parallel variance reduction in both populations. We note that 11 genes are not sufficient for a robust enrichment analysis despite several of them are relevant to digestion itself or feeding preference. *CG7025* (*FBgn0031930*), which is highly expressed in midgut (Leader et al., 2018). It was annotated as a digestive enzyme (Lemaitre & Miguel-Aliaga, 2013) and is part of the environment/microbiome-induced response in gut (Broderick, Buchon, & Lemaitre, 2014; Li et al., 2009; MacMillan et al., 2016). *CG1440* (*FBgn0030038*) is a peptidase involved in proteolysis (Larkin et al., 2021). It is expressed in digestive system (Larkin et al., 2021) and is related to the hedgehog signaling in response to dietary limitation (Çiçek et al., 2016). *Unc-119* (*FBgn0025549*), *Miga* (*FBgn0030037*), *CG12581* (*FBgn0037213*) and *CG32017* (*FBgn0052017*) are involved in olfactory memory formation (Walkinshaw et al., 2015), which may contribute to feeding preference.

## Discussion

### Change in gene expression variance is a selective response

Contrasting the gene expression of individual flies from ancestral and evolved populations, we studied the evolution of gene expression variance, a key parameter to understand adaptation which has not received sufficient attention in empirical studies. More than 100 genes experienced a significant change in variance during 100 generations of evolution in a novel hot environment. We consider this a conservative estimate because with a sample size of 20 individuals only rather pronounced variance shifts are significant. The analysis of more individuals probably would have identified even more genes with a significant change in variance. The significant enrichment of catabolism related GO categories indicates that the reduction in variance did not affect a random set of genes, as expected for stochastic changes occurring during 100 generations of evolution. While unlikely, it is conceivable that the enrichment for specific GO categories and tissues in a single population could be the outcome of genetic drift affecting a master regulator for these processes. Nevertheless, we observed that in two independent populations, not only the same GO categories and tissue enrichment was observed, but also more genes with a reduced variance were shared between the two populations than expected by chance. We conclude that the observed pattern of variance change reflects a shared selection response across independently evolved populations.

The lack of significant association between the genes with significantly reduced variance and the genes with significant mean changes indicates the operation of an independent stabilizing selection pressure reducing the variance of gene expression. Given that number of genes with significant differences in mean expression was much larger (>2500) than the number of genes with significant variance change (∼100), it might appear that selection on mean is stronger than selection on variance. Nevertheless, the power to detect changes in mean and variance differs and contributes to the discrepancy in the number of significant genes in the two categories. A larger sample size provides more power, but the difference in power to detect changes in mean and variance change in gene expression would persist.

### Potential selection pressures for the reduction in expression variance in gut

Based on the functional enrichment of catabolism related GO categories and enrichment for genes expressed in the gut, we considered different hypotheses that could explain the altered expression variance in the gut – microbiome, temperature and diet.

It is well-established that the microbiome has a pronounced effect on gene expression in the gut, but without a strong taxon-specific signal (Kokou et al., 2018). To investigate whether heterogeneity in microbiome complexity explains the evolution of gene expression variance, we used all remaining flies of the same common garden experiment from one evolved population and the reconstituted ancestral population (Supplementary file 1). The β-diversity, which quantifies the heterogeneity in microbiome complexity within a population, was very similar for evolved and ancestral populations (Supplementary figure 3 and Table 1). Despite the limitations of a very reduced sample size, our result is consistent with previous studies (Wong, Ng, & Douglas, 2011). The microbiome composition cannot explain the reduced variance as we observed high heterogeneity in composition among individuals from both the ancestral and evolved populations (Supplementary figure 3).

**Table 1.** Microbiome diversity in the reconstituted ancestral and hot-evolved population based on 16S-rRNA amplicon sequencing. ^a^ indicates the 95% confidence interval obtained from 100 bootstrap replicates.

|  | Ancestral | Evolved |
| --- | --- | --- |
| *α-diversity* | 33.84±0.90<sup>a</sup> | 27.60±0.57 |
| *β-diversity* | 1.42±0.04 | 1.37±0.03 |
| *γ-diversity* | 47.74±1.39 | 37.87±1.08 |

The experimental evolution study has been designed to study the impact of a shift in mean temperature, which remained constant across the entire experiment (Barghi et al., 2019). Natural populations experience substantial environmental heterogeneity – either seasonal or, due to microclimatic variation. As temperature modulates feeding preference and possibly digestion in *Drosophila* (Brankatschk et al., 2018), it is conceivable that some of the reduced expression variance might be explained by a more homogeneous temperature regime. Nevertheless, it is noteworthy that mean expression in several tissues are strongly affected by temperature – such as neural system and fat body (Hsu et al., 2020; Jakšić et al., 2020), but the involved genes do not experience a reduction in expression variance. In contrast, the reduction in gene expression variance in the gut implies specific selection pressure on the transcriptional variation of a subset of genes.

Alternatively, the reduced variance in midgut and digestion could also be caused by the unvarying diet in the laboratory. Natural *Drosophila* populations are feeding from different food sources in different microhabitats, that will result in a broader distribution of phenotypes in the ancestral population compared to laboratory evolved ones. This implies that such gene expression heterogeneity would be associated with fitness costs in a simple laboratory environment or specific expression patterns may be optimal on the laboratory food. Either scenario is consistent with polygenic adaptation driven by a narrower fitness function.

Given the experimental challenge to functionally validate putative selection pressures modifying expression variance, we performed computer simulations to illustrate our hypothesis that gene expression variance could be the direct target of selection during adaptation (Figure 4). We simulated a quantitative trait experiencing stabilizing selection over 200 generations and compared the dynamic of phenotypic variance with neutrality. Our results showed substantial decrease in phenotypic variance when stabilizing selection is imposed (Supplementary figure 4). This provides an illustrative support that the stabilizing selection caused by monotonic environment could alter the transcriptomic variation rapidly.

**Figure 4.**
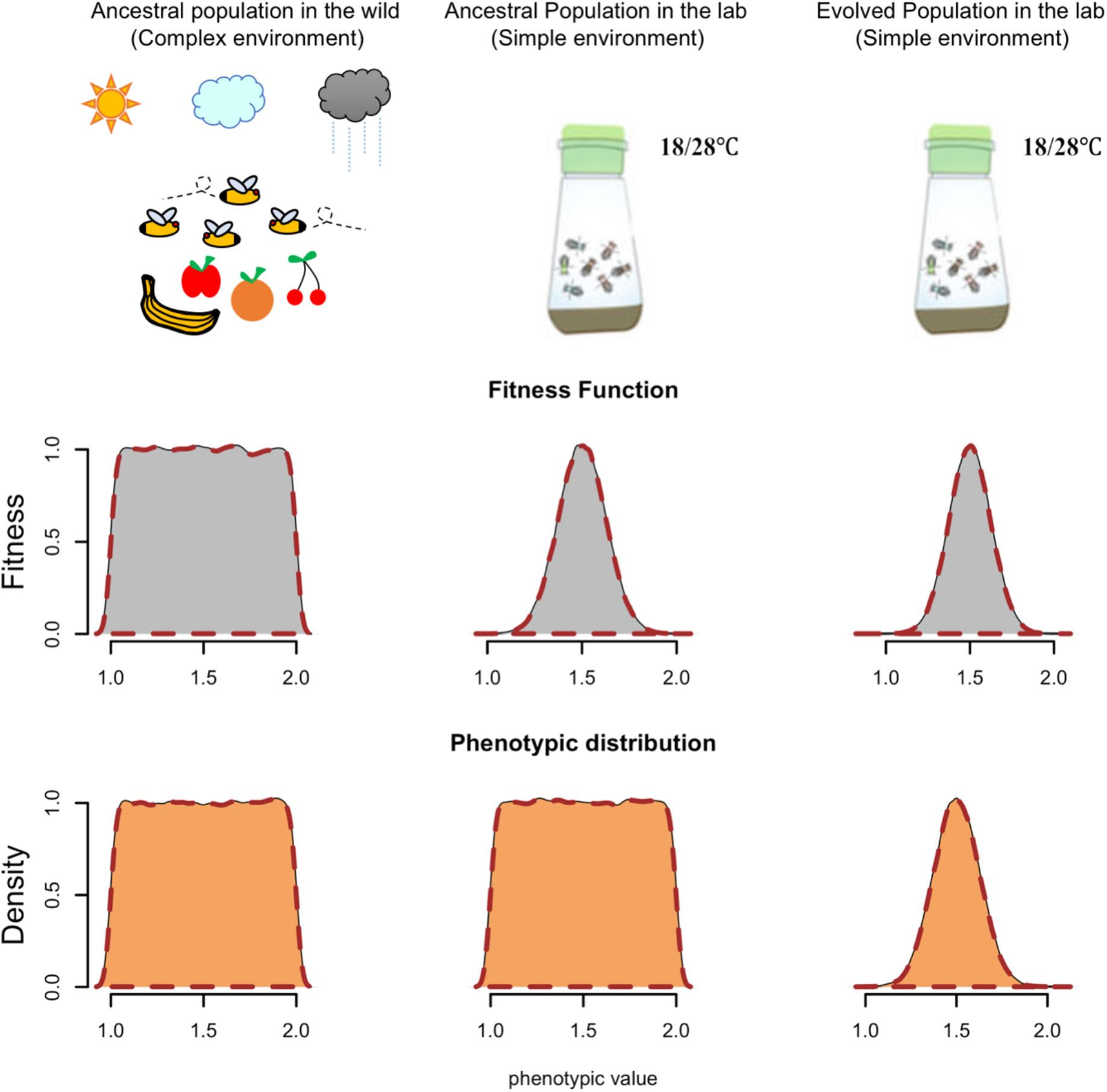
Hypothesis of a simpler lab environment selecting for decreased gene expression variance in the midgut. A proposed model for potential selection for decreased expression variance in midgut imposed by the drastic change in the environmental complexity. Food sources/fluctuation of temperature change dramatically when we bring these files from the wild into the lab. The distributions of fitness landscapes and expression value of the genes encoding digestive enzymes may change as the environmental shift. After 100 generations in a simpler environment, the genes encoding digestive enzymes decreased their expression variance.

Nevertheless, we acknowledge that canalization may have evolved and stabilizing selection could also act on the environmental component (i.e. expression noise of the genes) to reduce the phenotypic variance within population. Nevertheless, it is important to note that in this case canalization would act only on a subset of functionally related genes. Further work on canalization is required to determine the plausibility of this interpretation.

### Genetic redundancy and its regulatory basis

Another interesting observation was that both populations have different sets of genes with reduced variation (though the overlap is significantly more than random expectation), but both sets were enriched for genes expressed in the gut and the digestive enzymes (Supplementary file 6). Since we relied on a significance threshold to identify genes with a significant change in expression variance, the heterogeneity of the response in two independent evolved populations may (partly) be explained by stochastic effects reducing the power to detect the same genes in both replicates. To address this, we used a less stringent significance threshold (FDR of 0.1), but the heterogeneity in genes with significant expression differences between the two replicates persisted. Moreover, we show that resampling within the same replicate results in significantly more consistency than observed between the two populations (see material and method; Supplementary figure 5). This suggests that the stochastic effects cannot explain the heterogeneity observed between replicates. Rather, the heterogeneity also reflects the genetic redundancy during polygenic adaptation (Figure 5a; Barghi, Hermisson, & Schlötterer, 2020), which has been previously documented at the genomic level in the studied populations (Barghi et al., 2019). Our results are consistent with redundancy not only at the genomic but also at the gene expression level. If more contributing loci are segregating in the founder population than needed to reach the new trait optimum, the choice of redundant adaptive routes (variants/genes) is stochastic. This is particularly relevant during the early stage where drift affects the selection outcome (both for mean and variance) across replicate populations of moderate size (Bolnick, Barrett, Oke, Rennison, & Stuart, 2018; Langmüller & Schlötterer, 2020). With such initial stochasticity, the expression variance of digestion-related genes can be pushed in either direction. Henceforth, selection for variance reduction will favor genes for which drift acts synergistically with selection, leading to a heterogeneous outcome across populations (i.e. variance is reduced for different genes in the two replicates).

**Figure 5.**
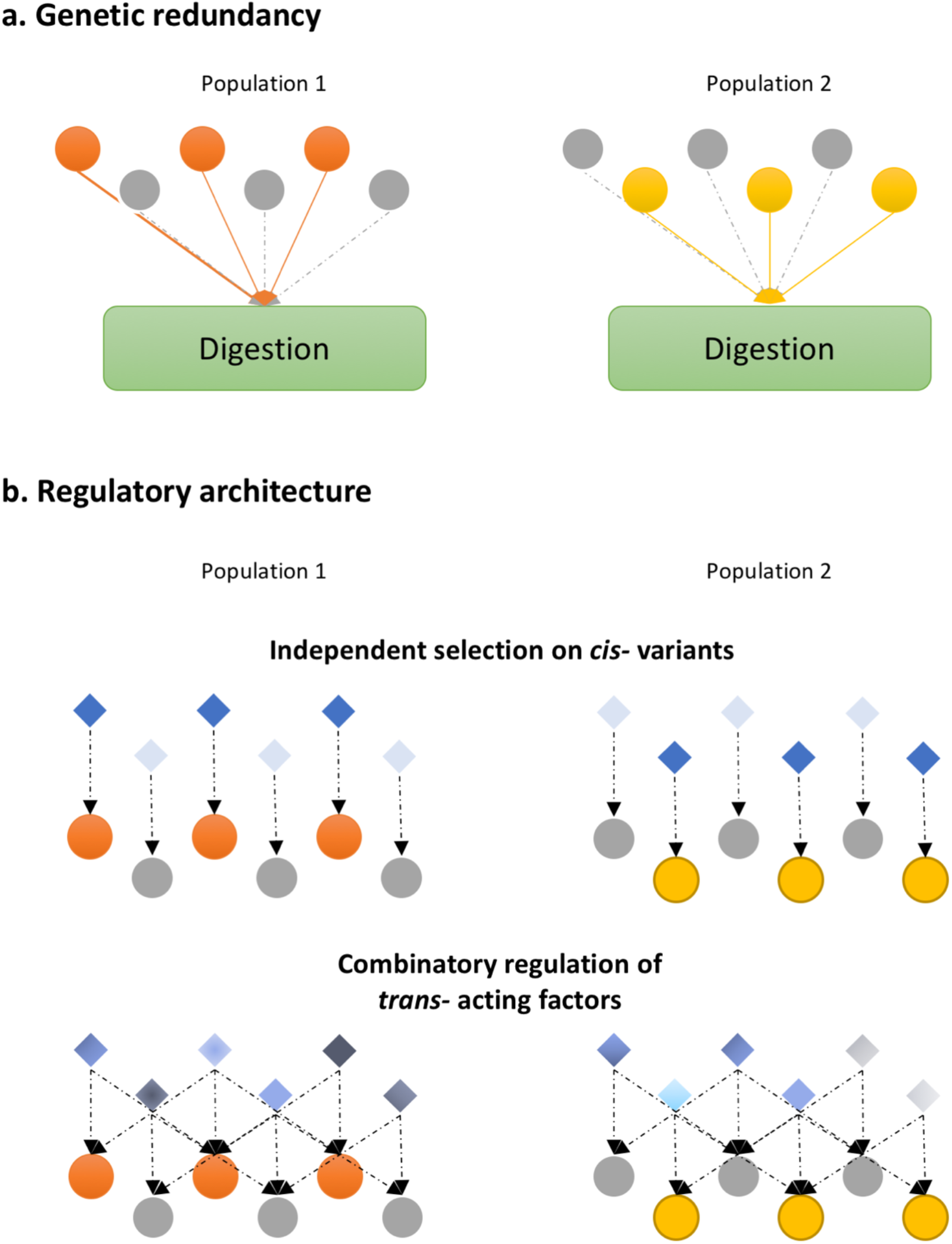
Schematic illustration of genetic redundancy at gene level with two possible regulatory architectures explaining the reduction in expression variance. **a**. Genetic redundancy: six genes contribute to digestion (higher-level phenotype) and the new phenotypic optimum could be reached by expression changes of three genes. Stochastic effects result in different gene sets (orange/yellow) responding to selection in the two populations. **b**. Two hypotheses about a regulatory architecture that allows for the rapid decrease in variance of digestion-related genes. Either selection acts independently on the *cis*-regulatory variants of each gene or combinatorial changes of several TFs reduce the expression variance.

While we discussed the genetic redundancy for genes involved in digestive function, the regulatory basis of the reduced variation is not yet clear. Gene expression can be regulated either in *cis* or in *trans. Cis*-regulation implies that independent regulatory variants are favored for each gene contributing to the selected phenotype (Figure 5b). It appears unlikely that each of the genes is independently targeted by selection. Rather, a more parsimonious explanation would be that several transcription factors (TFs) which cooperatively regulate these genes are the target of selection and reduced the expression variance of downstream genes. We explored this hypothesis and searched for *trans*-acting TF binding sites shared among genes with decreased expression variance and high expression in the midgut. We identified 18 and 8 TFs in the two populations, some of which evolved their mean expression (Supplementary file 7), but none evolved a significant change in expression variance. The lack of significant variance evolution in these candidate targets of selection suggests a more complex regulation of transcriptional variance. We consider it highly likely that the expression of each redundant gene is in turn regulated by several trans-acting factors – providing a second layer of possible genetic redundancy (Figure 5b). We attempted to incorporate previously documented genomic signatures of selection (Barghi et al., 2019) into the regulatory basis of the evolution of gene expression variance. Interestingly, we found that ∼50% of the candidate genes/TFs covered at least a significant SNP 5kb up-/down-stream the genic regions. However, given the large number of selection targets, this number is not significantly more than random expectation (Permutation test, p-value > 0.05). Hence, no conclusive inference on the complex regulatory basis of redundant gene expression can be achieved from the genomic data. Clearly, more work is needed to study the regulatory architecture of genetic redundancy, but the experimental framework introduced here provides a starting point.

## Concluding remarks

Previous studies on adaptive phenotypic evolution mainly focused on “population means”, to explain adaptation to different environments. Nevertheless, also selection on “phenotypic variance within a population” (e.g.: stabilizing selection, disruptive selection…) has been studied, but the possibility that environmental changes affect the strength of stabilizing selection has not yet been sufficiently explored. In a recent commentary (Harpak & Przeworski, 2021) highlighted that environmentally triggered changes in the strength of stabilizing selection may have important implications for the understanding of polygenic traits. To our knowledge, our study provides the first empirical evidence that the shift from a complex natural environment to a simple laboratory environment could result in drastic changes in the expression variance of certain genes during adaptation. This has important consequences for future research on adaptive phenotypic evolution: In addition to searching for changes in mean phenotype as a response to selection, it is also important to consider that phenotypic variance may be subject to selection and can contribute to our understanding of adaptation processes in natural and experimental populations.

## Acknowledgments

We especially thank Viola Nolte for preparing all RNA-Seq and 16s-rRNA libraries, and supervising the maintenance of the evolution experiment. We thank all member of the Institut für Populationsgenetik for discussion. Ana Marija Jakšić, Neda Barghi, François Mallard and Kathrin Otte contributed to the common garden experiment. Illumina sequencing was performed at the VBCF NGS Unit (www.vbcf.ac.at). This work was support by the Austrian Science Funds (FWF, W1225) and the European Research Council (ERC, ArchAdapt).

## Author contribution

W.Y.L and C.S. conceived the study. W.Y.L performed the data analysis. W.Y.L. and C.S. wrote the manuscript.

## Data accessibility statement

All sequencing data are available from the European Nucleotide Archive (ENA) under the accession number PRJEB37011. Original data and scripts for the analysis are available on the GitHub repository of this study (https://github.com/cloudweather34/variance).

## Correspondence and requests for materials

should be addressed to C.S.

---

**Supplementary figure 1.**
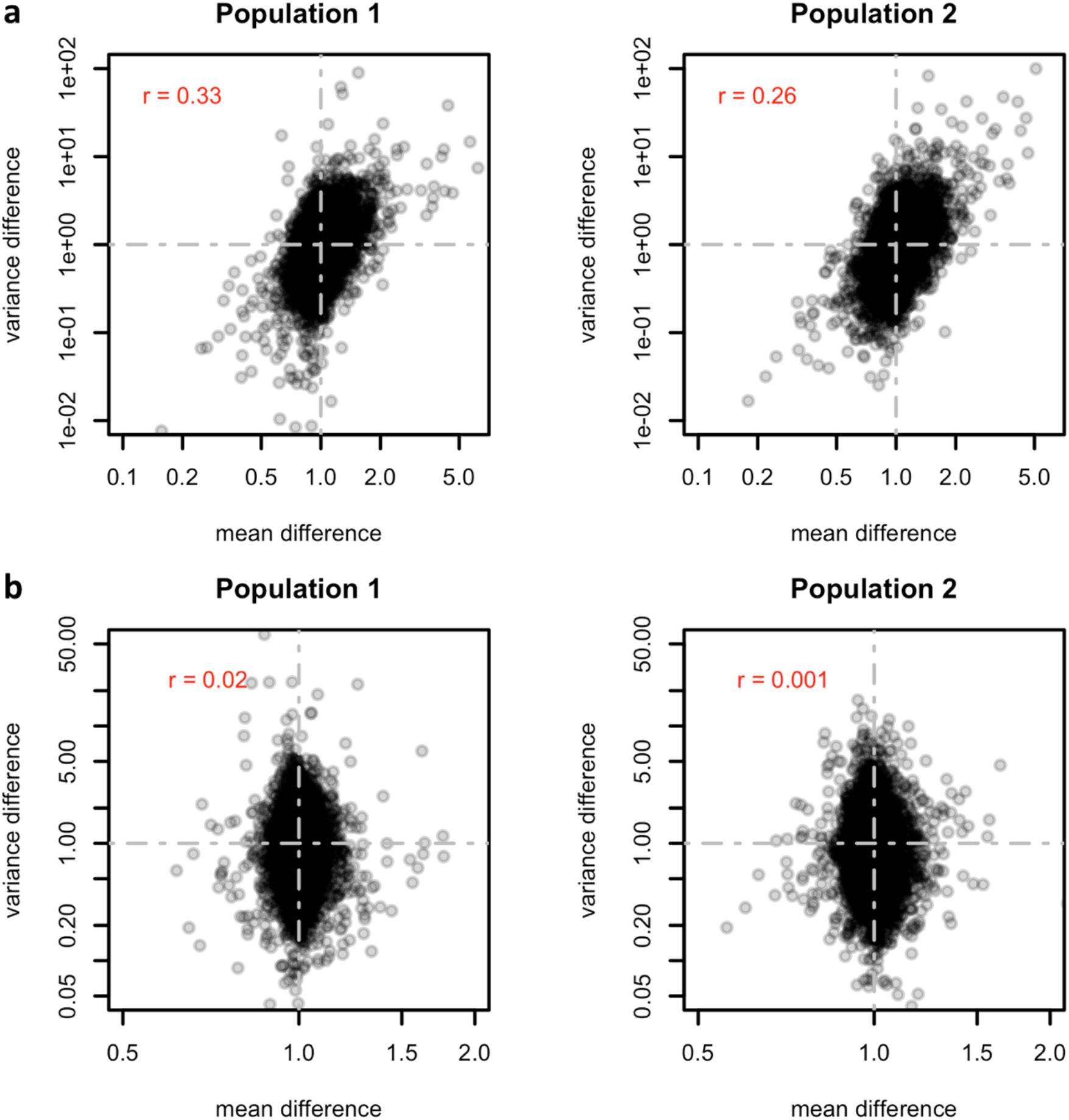
Log-transformation eliminates the positive relationship between the changes in mean and variance of gene expression. In each panel, the difference in mean expression 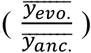 and the difference in variance 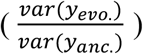 before **(A)** and after **(B)** the natural log-transformation of each gene were visualized. The systematic positive correlation in the differences due to the positive mean-variance dependency of negative binomial distribution is removed by the log-transformation on gene expression level.

**Supplementary figure 2.**
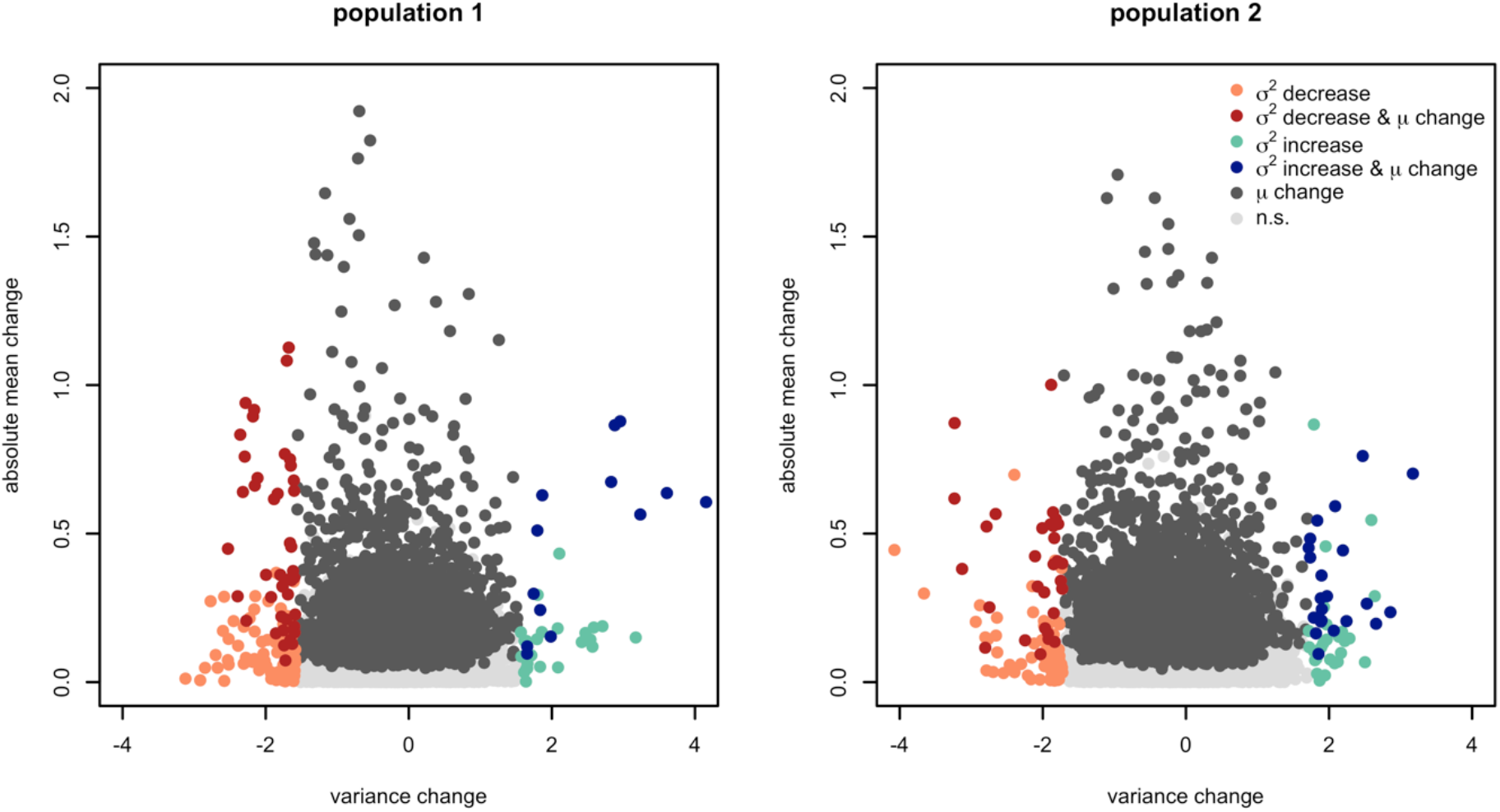
No correlation between the evolutionary response of variance and mean: evidence from genes which changed significantly in mean or variance. Each gene is plotted for the two populations with the change in variance 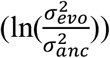 on the X-axis and absolute mean change (|*μ*_*evo*_ − *μ*_*anc*_|) on the Y-axis. Genes with significant changes in variance (depicted in color) fall into both categories-with and without significant changes in mean expression. Only a negligible correlation was detected in a correlation analysis of the changes in variance and mean experiment in the 1^st^ (left panel; rho = 0.047) and 2^nd^ (right panel; rho = -0.001) population among the genes with significantly altered means or variances. The negligible correlation suggests an independent selection response for mean and variance.

**Supplementary figure 3.**
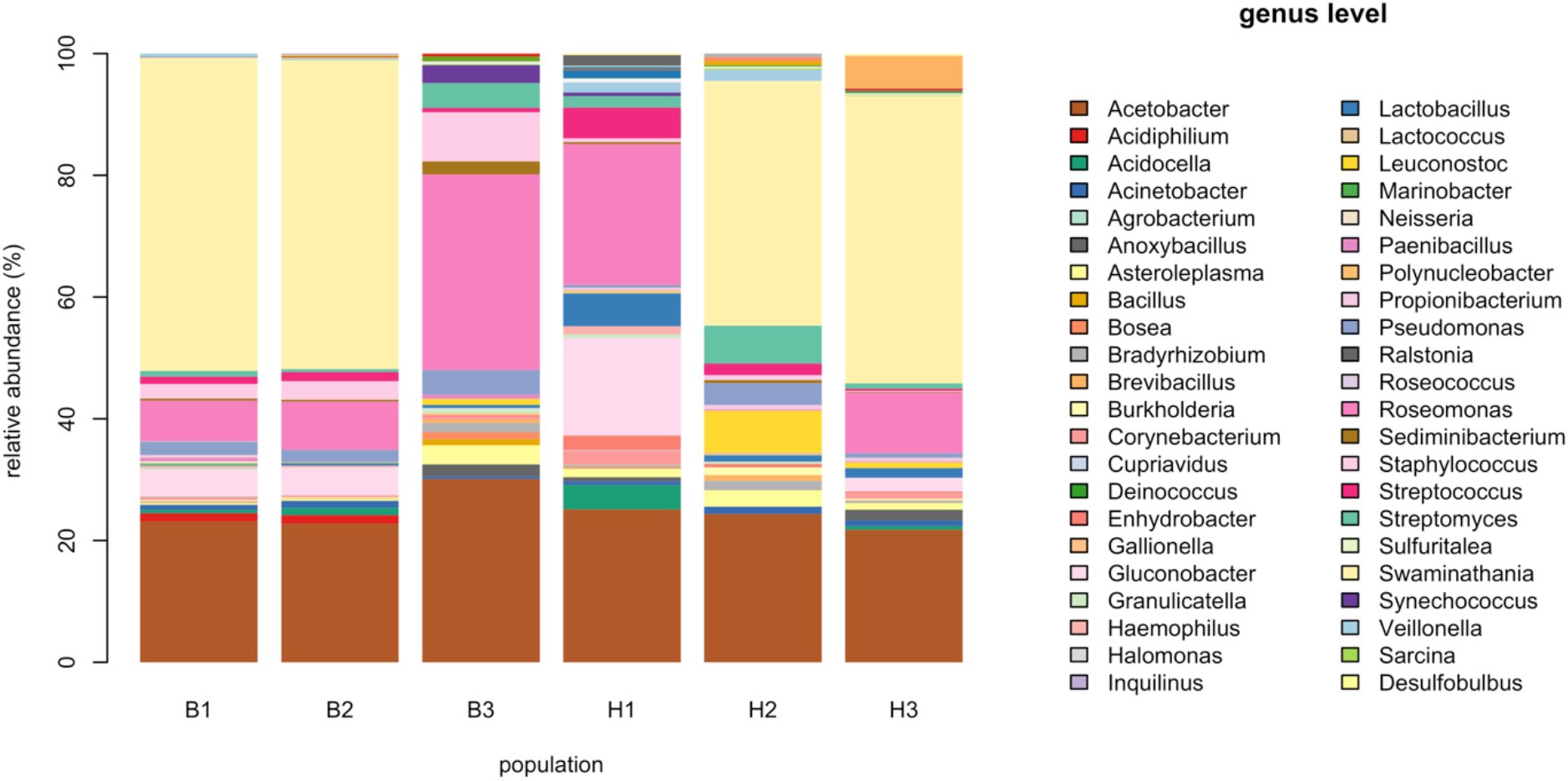
Microbiome composition in ancestral and evolved flies. Microbiome composition on the genus level for three individuals from the ancestral population (A1-A3) and five individuals from a hot-evolved population (H1-H3).

**Supplementary figure 4.**
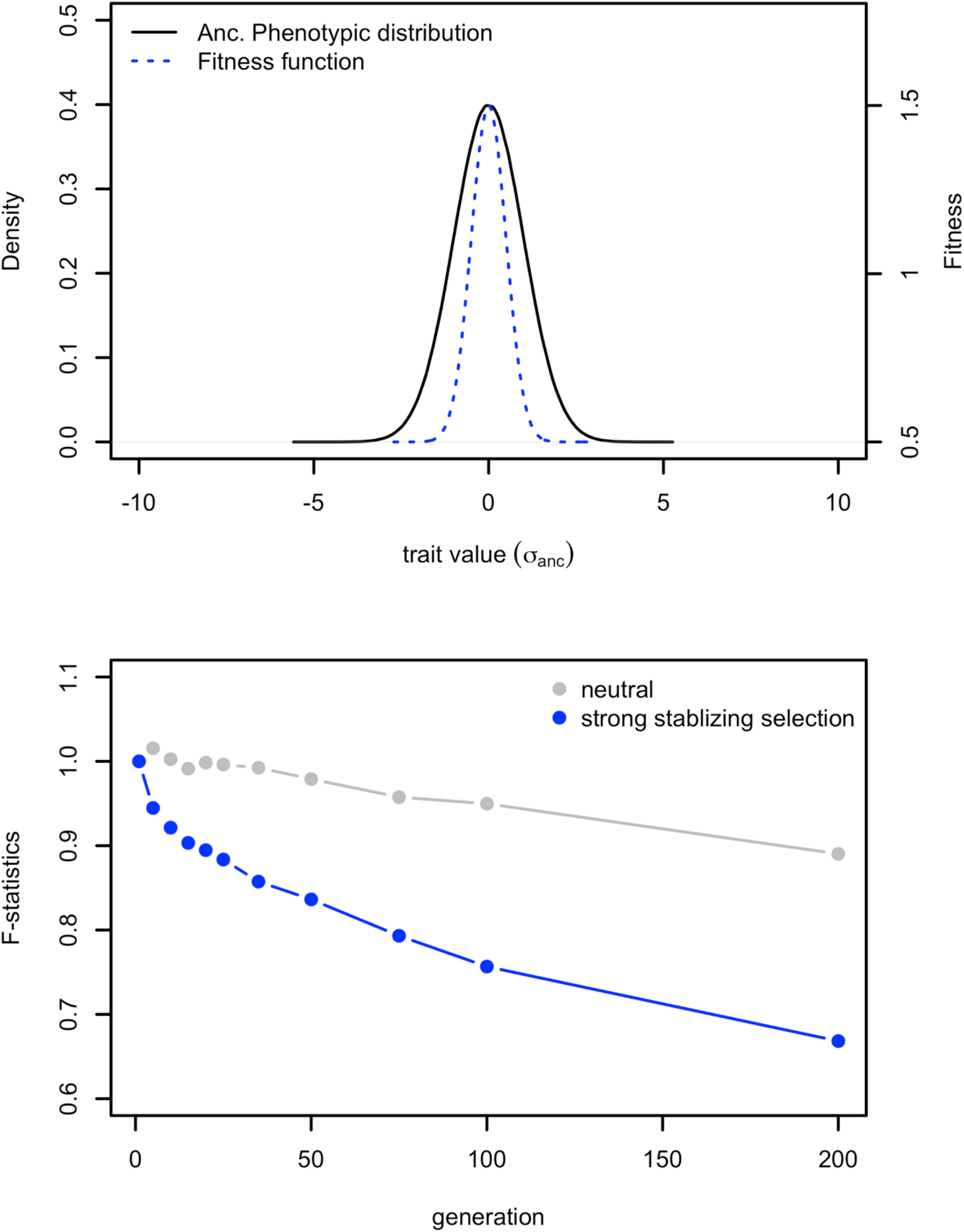
Reduction in variance by strong stabilizing selection. **a**. Computer simulations of a scenario where the shift to a simpler environment results in stronger stabilizing selection. The ancestral phenotypic distribution of quantitative trait under stabilizing selection before the population was introduced to the simple environment (black). The fitness function after the habitat shift is shown in blue. The variance of the fitness function is set to 0.5 standard deviation of the ancestral trait distribution. **b**. The changes in phenotypic variance under strong stabilizing selection (blue) and neutrality (grey). The change in phenotypic variance (F) is calculated as the ratio between the evolved and ancestral phenotypic variance at each generation 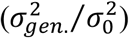 for each scenario. For each scenario, 1000 simulation runs were performed.

**Supplementary figure 5.**
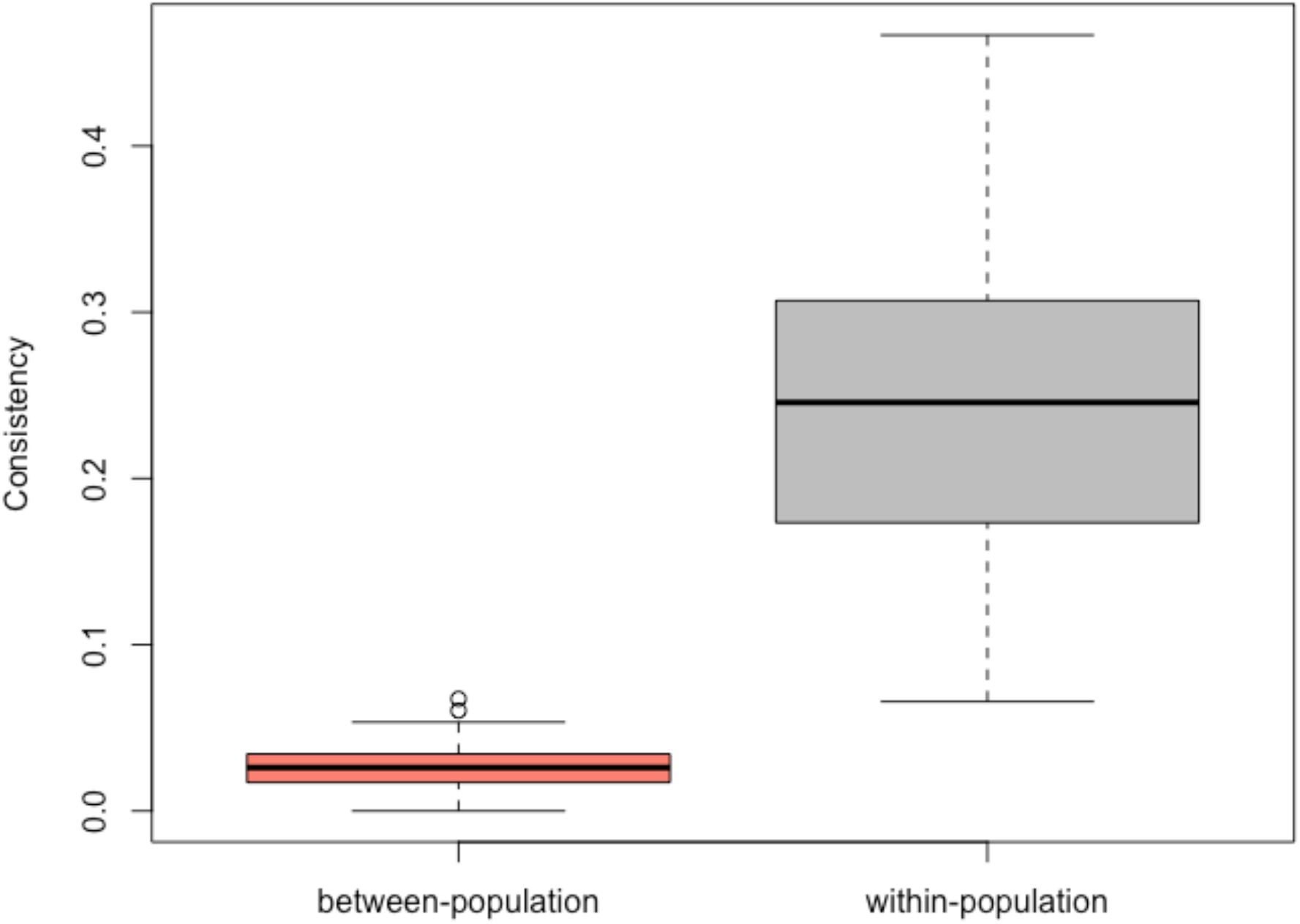
Heterogeneous selection response between the two populations cannot be explained by sampling variation. To demonstrate that the observed heterogeneity between two evolved populations is not due to the stochastic effect, within the first comparison (ancestral population 1 vs. evolved population 1), we subsampled two sets of 15 individuals from the ancestral and evolved population and performed tests for variance (contrasting the evolved and ancestral samples) on this two sets independently. We then estimated the consistency between tests using the Jaccard Index. This process is repeated 100 times, generating the null distribution of Jaccard Indices considering sole stochastic effects within an evolutionary replicate (grey). Similar sampling scheme for 15 individuals was applied to the comparison between the two populations 100 times. The observed Jaccard indices between two populations are shown in red box and falls clearly outside of the null distribution. The significant higher consistency in the resampling analysis within the same replicate (Wilcoxon’s test, W = 9999, p-value < 2.2 × 10^−16^) suggests that the stochastic effects along cannot explain the heterogeneity observed between replicates.

